# IgG surface mobility promotes antibody dependent cellular phagocytosis by Syk and Arp2/3 mediated reorganization of Fcγ receptors in macrophages

**DOI:** 10.1101/2021.07.12.451665

**Authors:** Seongwan Jo, Nicholas M. Cronin, Ni Putu Dewi Nurmalasari, Jason G. Kerkvliet, Elizabeth M. Bailey, Robert B. Anderson, Brandon L. Scott, Adam D. Hoppe

## Abstract

By visualizing the movements of Rituximab during Antibody dependent cellular phagocytosis (ADCP) of B lymphoma cells by macrophages, we found that Fcγ receptors (FcγR) on the macrophage surface microcluster, recruit Syk and undergro large-scale reorganization at the phagocytic synapse prior to and during engulfment of the target cell. Given these dramatic rearrangements, we analyzed how the surface mobility of Rituximab contributes to FcγR signal amplification and ADCP efficiency. Depolymerization of the target cell actin cytoskeleton resulted in free diffusion of Rituximab docked to CD20, enhanced microcluster reorganization, Syk recruitment and ADCP. Conversely, immobilization of Rituximab by chemical fixation impaired microcluster formation and diminished Syk recruitment and ADCP. In macrophages lacking Syk, Rituximab accumulated at the base of the phagosome and were trogocytosed, consistent with Syk kinase activity being necessary to trigger redistribution of Rituximab-FcγR during engulfment and to prevent antigenic modulation of the target. Total internal reflection fluorescence analysis of FcγR-IgG on fluid supported lipid bilayers revealed a membrane topography displaying inward reaching leading edges and protruding contact sites reminiscent of podosomes. This topography was distinct from the closely apposed macrophage/target membranes observed during engagement of IgG displayed on immobile supported lipid bilayers. The organization of this contact, pseudopod extension and the rearrangement of microclusters depended critically on Arp 2/3. Thus, Syk and Arp2/3 coordinate actin rearrangements and FcγR-IgG complexes that were of previously unrecognized complexity for the clearance of cells displaying surface-mobile antigens.

**Significance Statement:** ADCP is an important effector mechanism for the removal of malignant, immunologically aberrant, and infected cells during treatment with therapeutic antibodies or adaptive immune responses. Most transmembrane protein antigens are mobile with transient confinement from the actin of the target cell. This work demonstrates that macrophage forces overcome these confinements to rearrange FcγR-IgG-antigen complexes before and during ADCP. Thus, new paradigms are needed as ADCP has largely been studied using model target particles that display immobile antigens. Moreover, we found that the mobility of the therapeutic antibody, Rituximab, on the surface of B lymphoma cells foretells ADCP efficacy, with lower densities of IgG mediating ADCP on increasingly mobile antigens.

## Introduction

Therapeutic antibodies are a rapidly growing class of drugs for treating rheumatological, viral, and malignant conditions (1–3). Nearly all are of class IgG owing to long circulatory half-lives, broad tissue distribution, and the ability to activate potent immune effector functions on cells bearing Fcγ receptors (FcγR)(4–6). A growing body of evidence indicates that antibody dependent cellular phagocytosis (ADCP) by macrophages and monocytes is an important or essential effector mechanism, especially for the destruction of cells of lymphocytic origin (4, 7–11). Some of the most successful therapeutic antibodies that mediate ADCP killing of lymphocytes include anti-CD20 (Rituximab) and its derivatives, and anti-CD52 (Alemtuzumab) (2, 8, 12, 13). Conversely, ineffective ADCP can cause trogocytosis, in which FcγR bearing effectors remove small bites of the target cell’s antigen and plasma membrane leading to antigenic modulation (Rituximab depletion of CD20 from B cell lymphomas) or killing of large target cells (Cetuximab on breast cancer cells) (14–17). Thus, there is a pressing need to understand the cellular and molecular mechanisms by which FcγR ligation to target-bound IgG orchestrates effective ADCP, and how the antigen on the target cell influences this process to enable the development of improved therapeutic antibodies and adjuvants.

Macrophages, monocytes, and dendritic cells express all FcγRs to varying degrees and are resident in nearly all tissues of the body, making them key effectors of IgG-mediated responses and antigen presentation (4, 8, 11, 18–20). FcγR function on macrophages contributes to therapeutic antibody anti-tumor activity, provided that inhibitory signals originating from the microenvironment are overcome, including the use of STING agonists to promote ADCP and CD47/SIRPα blocking antibodies to alleviate innate inhibitory signaling in tumor microenvironments (8, 20–23). Recently, the concept of engineering macrophages with chimeric antigen receptors, analogous to CAR-T cells, as an alternative cell-based therapeutic strategy has emerged (24), further highlighting the need to understand the molecular mechanisms regulating macrophage ADCP. Our understanding of ADCP and the mechanisms underpinning its efficacy and failure are shaped by the two historical models of phagocytosis known as the ‘zipper’ and the ‘trigger’(25–27). The zipper model posits that FcγRs must sequentially engage IgG on the target surface as actin polymerization extends membrane over the target, like the closing of a zipper (25). In contrast, the trigger model posits that if sufficient FcγRs bind IgG, an activating signal will trigger large-scale cytoskeletal rearrangements to engulf the target (27). In seminal work from the Silverstein lab, the zipper model prevailed. Specifically, they observed that when IgG was ‘capped’ onto one side of the lymphocyte surface macrophages failed to engulf them, rather leading to trogocytosis of the IgG with diffuse IgG distributions associated with successful ADCP(25). Subsequently, work suggested that the targets cells that rearranged CD20-Rituximab to their uropod, creating asymmetrical arrangements that lead to failed ADCP, and trogocytosis by RAW264.7 cells (28). However, extensive reorganization of IgG-FcγR by macrophages on synthetic membranes (29, 30) and on cancer cells anti-CD19 (31) has been observed. Moreover, IgG antibodies vary widely in their ability to mediate therapeutic responses (32, 33). Thus, understanding how the organization of FcγR-IgG will provide critical information for the rational design of next-generation therapeutic antibodies including establishing criteria for antigen selection, optimal antibody Fc structure, and complementary adjuvant therapies to promote ADCP.

Here, we have addressed how cellular rearrangements of Rituximab influence FcγRs signaling at the macrophage/target cell interface during ADCP. Our findings indicate that the surface mobility of Rituximab has a profound effect on successful ADCP, with increased surface mobility leading to increased Syk recruitment and improved likelihood of successful ADCP. Moreover, we find that Syk and Arp2/3 work together to organize IgG-FcγR microclusters prior to and during engulfment and limit trogocytic clearance of IgG from the target cell.

## Results

### Rituximab is microclustered and pools at the base of the phagosome where it amplifies FcγR signaling prior to being disbursed during target cell engulfment

Given our prior findings that IgG presented on fluid SLBs is rearranged by macrophage FcγRs in a process reminiscent of TCR and BCR rearrangements, we hypothesized that similar rearrangements occur during ADCP (30). To address this question in a clinically relevant context, we sought to determine the movements of Rituximab (RTX) on the surface of B cell lymphoma targets during ADCP. Confocal microscopy of opsonized WIL2-S cell ADCP by FLMs revealed that CD20-docked RTX-AF647 was rearranged into microclusters upon contact with the macrophage (Fig. 1, Video 1). Moreover, a previously validated Syk-mScarlet construct (34), was recruited to these microclusters, thus demonstrating a high degree of phosphorylation. During engulfment, the RTX-FcγR-Syk microclusters were redistributed over the surface of the WIL2-S target as the enveloping pseudopod advanced, ultimately producing a phagosome containing the WIL2-S target with an approximately uniform distribution of RTX-FcγR-Syk microclusters forming the connection between phagosome and target membranes (Fig. 1A). To gain a more complete three-dimensional view of this process, we applied lattice light sheet microscopy (LLSM), capable of capturing 3D imaging volumes with minimal photoxicity at high imaging frame rates (35). LLSM imaging revealed that RTX-AF647 present on the surface of WIL2-S cells appeared somewhat punctate owing to the filipodia present on the WIL2-S surface, rather than a pre-clustering of the RTX. Upon contact with the macrophage, the RTX-AF647 formed microclusters that rapidly recruited Syk-mScarlet (Fig. 1B, Video 2). As the phagocytic cup began to form, the microclusters assembled into larger patches as they streamed toward the base of the phagosome (initial contact site), suggesting a coupling between the macrophage FcγRs and the retrograde flow of the actin cytoskeleton. Within 2-3 min post contact, membrane ruffles, not attached to the target began to emerge, radiating in a flower-like pattern around the engaged target as RTX microclusters continued to accumulate into larger patches at the base of the forming phagosome and recruit Syk-mScarlet. Extension of the pseudopod followed, during which many microclusters were propelled forward over the target, resulting in a nearly uniform distribution of RTX-Syk microclusters over the internalized target (Fig 1B, Video 2). These behaviors were consistent over 12 confocal and 3 LLSM movies. Taken together, these data indicate that the macrophage cytoskeleton drives rearrangement of RTX docked to FcγRs during ADCP in two phases: 1) initial microclustering followed by accumulation at the base of the phagocytic cup, and 2) an engulfment phase in which these microclusters redistribute along the advancing pseudopod. This finding contrasts with prior observations using macrophage-like RAW264.7 cells, which exclusively perform trogocytosis of RTX when engaging Raji cells (28). Here, most interactions resulted in complete ADCP of the WIL2-S cell with a small portion of the RTX being packaged into small vesicles that separated from the WIL2-S cell prior to and during engulfment (Fig. 1B, Video 2).

**Figure 1.**
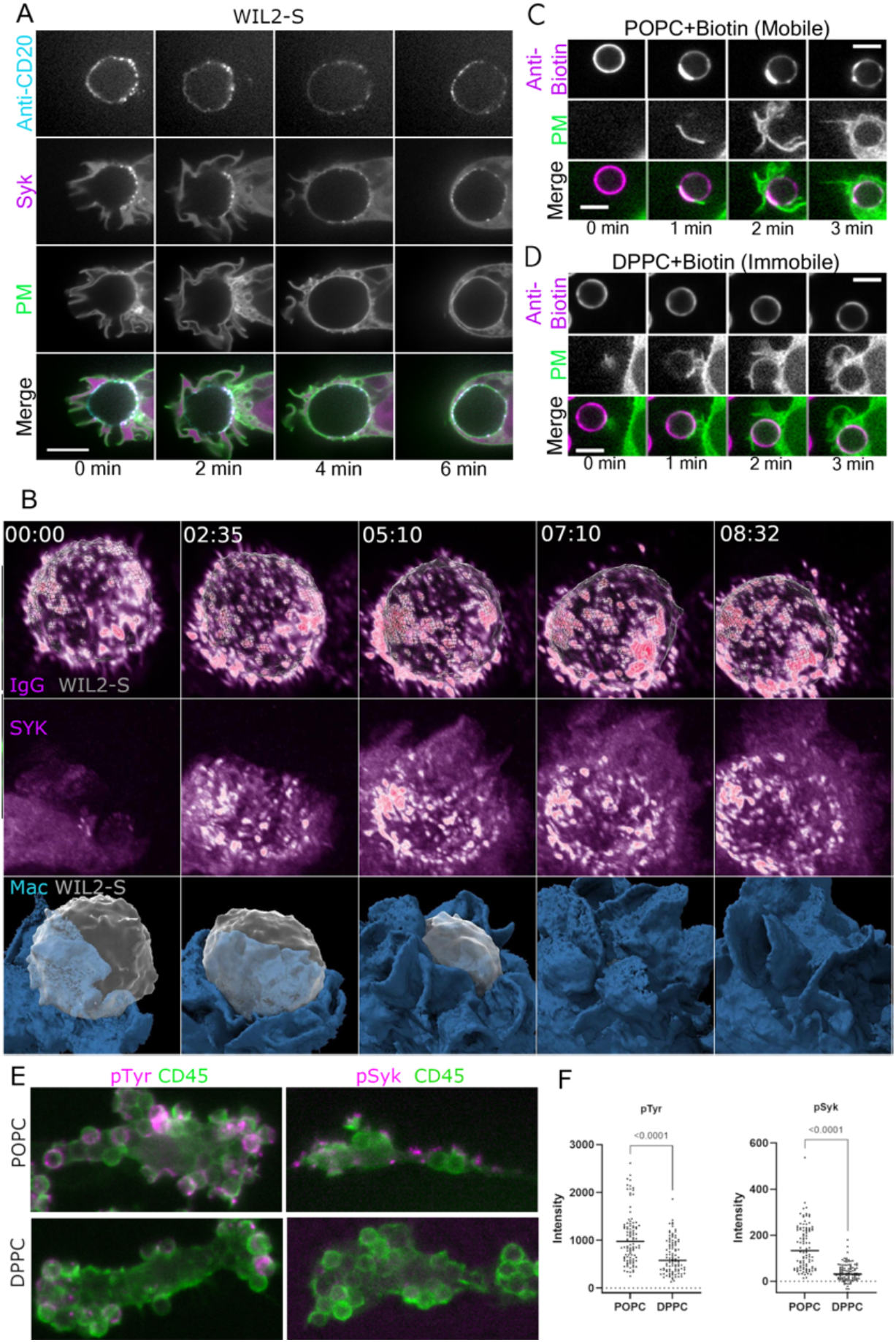
Spatial reorganization of RTX and enhanced FcgR signaling on mobile IgG surfaces during ADCP. (**A**) Confocal imaging of macrophage-induced microclustering and rearrangement of RTX-AF647 on WIL2-S cells. Syk-mScarlet and mNG-Mem were stably expressed in FLM macrophages. During engulfment, RTX-AF647 microclusters docked with Syk-mScarlet redistributed over the target surface (scale bar = 10 µm). (**B**) LLSM imaging of ADCP of WIL2-S cells labeled with calcein violet and RTX-CF647 by an FLM expressing mNG-Mem and Syk-mScarlet, illustrated dynamic formation of microclusters accumulating in a ring at the base of the target-macrophage interface (5:10 min). As the engulfment proceeded, RTX microclusters recruited Syk-mScarlet and redistributed over the surface of the WIL2-S cell (7:10 and 8:32 min, image dimension = 20 µm x 20 µm). Anti-biotin mIgG2a-AF647 displayed on SLBs on silica beads was rearranged by PKH67-labeled (PM) FLMs into a patch at the phagosome base and then redistributed over the bead when fluid POPC (**C**) but not for gel-phase/immobile DPPC SLBs (**D**) (scale bar = 5 µm). **(E)** Phosphotyrosine and phospho-Syk (pTyr525/526) immunofluorescence (magenta, counterstained with anti-CD45, green) in BMDMs 12 min after being fed silica beads displaying similar densities of IgG on mobile (POPC) or immobile (DPPC) SLBs. (**F**) Quantification of individual forming phagosomes indicates elevated pTyr and pSyk. Student’s t-test comparison.

To determine if the forces responsible for reorganizing the RTX were generated by the macrophage, we imaged the redistribution of IgG on fluid and immobile supported lipid bilayers on silica beads (SLB-SB). The mobility of SB-SLBs was validated by FRAP followed by TIRF imaging (Supplement Fig. 1). Upon contact with fluid POPC SLB-SBs, anti-biotin IgG pooled at the initial contact site indicating it was being captured by FcγRs (Fig. 1C). Engulfment of the SLB-SB was coincident with redistribution of the anti-biotin-IgG-AF647 over the bead surface, recapitulating the overall movements observed for RTX on WIL2-S targets (Fig. 1B,C). When SLB-SBs were made from gel-phase/immobile DPPC, little or no change in the distribution of anti-biotin-IgG was observed (Fig. 1D). Thus, we conclude that the macrophage generates sufficient forces to capture and reorganize IgG on fluid bilayers.

We used our SLB-SB system to determine if microclustering of mobile IgG promotes FcγRs signal amplification as has been reported for BCRs and TCRs (36–38). Indeed, immunostaining of BMMs trapped during phagocytic cup formation (∼12 min after addition of SLB-SB targets) demonstrated that tyrosine phosphorylation (pTyr) and Syk phosphorylation (pSyk) was significantly elevated when similar densities of IgG (Supplemental Fig. 1) were presented on fluid SLB-SBs composed of POPC vs. on immobile SLB-SBs composed of DPPC (Fig. 1E,F). Taken together, these data indicate that ADCP takes place in three phases: 1) engaged FcγRs are initially microclustered upon contact with the target and binding to IgG 2) signal amplification during which these microclusters recruit Syk and phosphorylate other proximal signaling machinery leading to cup initiation and ligation of additional FcγRs and pooling together of microclusters into patches that continue to recruit Syk and 3) engulfment, during which organized redistribution of the microclusters at occurs at the advancing pseudopod edge. Thus, if given permissive antigen mobility, macrophage forces will drive substantial rearrangements of target associated IgG.

### The mobility of RTX on WIL2-S cells promotes ADCP

To determine if the degree of antibody mobility affected ADCP the efficiency of ADCP, we manipulated the cytoskeleton of WIL2-S cells to liberate or restrict the mobility of CD20 (39). The phagocytic index (number of target cells internalized per macrophage) was quantified as a function of RTX density by high content microscopy (Fig. 2). Treatment of WIL2-S cells with the actin depolymerizing drug, Lat B, imparted free diffusion to RTX and increased the ADCP response relative to non-treated WIL2-S cells (Fig. 2). In contrast, stabilization of the WIL2-S cytoskeleton and surface topography with the combination of Jasplakinolide and Blebbistatin (Jas/Bleb) (40) had no discernable effect on the mobile fraction of CD20 or on ADCP (Fig. 2), consistent with the WIL2-S cytoskeleton confining CD20 and not playing an active role in RTX organization. In contrast, fixation of WIL2-S cells with paraformaldehyde (PFA), to immobilize CD20 resulted in a higher RTX density needed to promote ADCP (Fig. 2). Notably, FRAP measurements of RTX-AF647 demonstrated limited diffusion for untreated cells (mobile fraction < 10%, Fig 2C). However, the forces created by engagement with macrophage FcγR leads to its rearrangement on the cell surface, (Fig. 1) suggesting cytoskeletal pickets within the target cell that create CD20 diffusion barriers that can be hopped with small forces (41–43). To control for changes in the cell surface abundance of CD20 during these treatments, RTX density was measured by flow cytometry and used to define the IgG-surface density (Fig. 2B, Supplemental Fig. 2). Thus, the mobility of RTX was a strong predictor of the potency of RTX in promoting ADCP.

**Figure 2.**
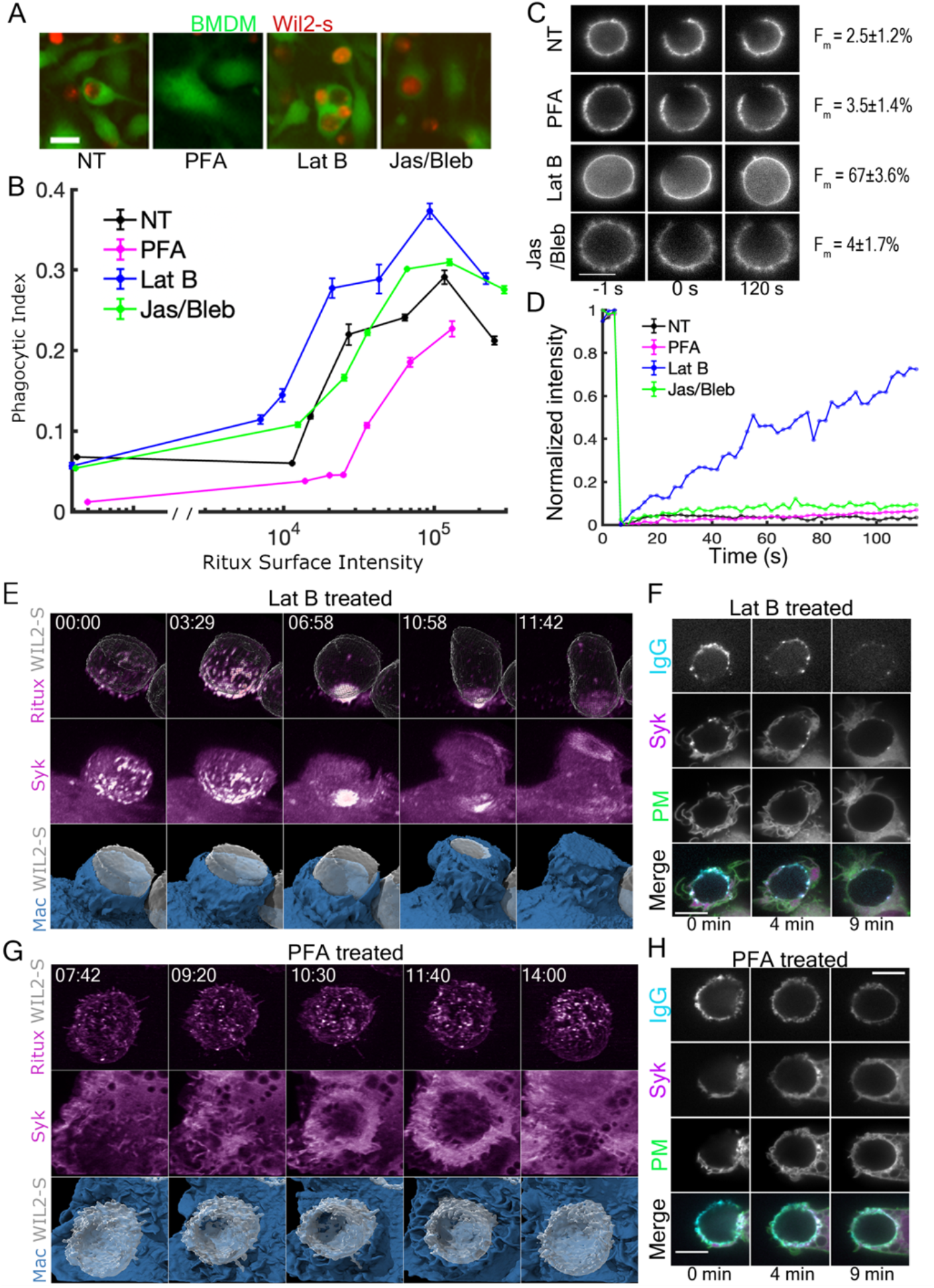
Enhanced ADCP and IgG/Syk microcluster rearrangement with increasing CD20 surface mobility. (**A**) High content imaging of BMDMs (green) and internalized WIL2-S cells (pHrodoRed) opsonized with RTX following manipulation of actin (NT, PFA, LatB and Jas/Bleb, scale bar = 10µm). (**B**) Quantified phagocytic index for measured RTX densities on WIL2-S cells that were untreated or pretreated with LatB, Jas/Bleb or PFA (error bars are SEM, n = 2 experiments). (**C** and **D**) FRAP images and normalized recovery traces of anti-RTX-AF647 on single WIL2-S cells treated with the indicated actin disrupting drugs and the average mobile fraction F_M_, for 10 cells from each condition (error is std. dev., scale bar = 10 µm). (**E and G**) LLSM of calcein violet labeled WIL2-S cells opsonized with RTX-CF647 and pretreated with Lat B during complete ADCP (**E**) or treated with PFA during failed ADCP (**G**) by FLMs expressing mNG-Mem (PM) and Syk-mScarlet. For the Lat B treatment, RTX-CF647 microclusters accumulated into a large patch at the base of WIL2-S-FLM contact site where Syk-mScarlet was recruited. Redistribution of this patch led to near uniformity during engulfment (11:42 min). Limited rearrangement of RTX-CF647 into small clusters occurred on PFA treated WIL2-S cells (image dimension = 21 µm x 21 µm for E and 25 µm x 25 µm for G). (**F and H**) Confocal imaging during complete ADCP of WIL2-S cells treated with LatB (**F**) or PFA (**H**, scale bar = 10 µm).

Given the enhancement in ADCP produced by LatB treatment of the WIL2-S cell, we examined the movements of RTX by LLSM and confocal imaging. These movies revealed that that LatB treatment of the WIL2-S cell resulting in free diffusion of RTX-CD20 (Fig 2C,D) produced dramatic microclustering, and pooling of RTX at the base of phagosome followed by extensive redistribution during engulfment (Fig. 2E,F), when compared with non-treated WIL2-S targets (Fig. 1A,B). Thus, the forces generated by macrophage FcγRs dramatically reorganized RTX and produced intense recruitment of Syk-mScarlet at the base of phagosome that was redistributed over the target during engulfment (Fig 2E and F). For target cells treated with LatB, the degree of RTX pooling at the base of the phagosome was dramatic and closely resembled the that of IgG presented on fluid SLB-SBs (compare Fig. 2E with Fig. 1C). Conversely, imaging ADCP of PFA-treated WIL2-S cells demonstrated minimal rearrangement of RTX (Fig. 2 G-H) and minimal Syk-mScarlet recruitment in the LLSM example of failed ADCP (Fig. 2G), and low levels of Syk-mScarlet recruitment in the confocal imaging example of successful ADCP of a PFA treated WIL2-S cell (Fig. 2H). In comparing these two examples, with the movements of RTX on unperturbed WIL2-S cells, it is clear that forces from the macrophage FcγRs overcome CD20 corralling by the target cell actin cytoskeleton and drive its rearrangement (Fig 2C,D). Together, these findings indicate that microclustering, followed by pooling of RTX-FcγR complexes at the base of the phagosome is driven by forces originating from the macrophage and contributes to FcγR signaling, Syk recruitment, and successful ADCP.

### IgG surface mobility shapes the topography of the macrophage-target interface

The nanoscale topography of the macrophage plasma membrane is thought to be important for shaping signal amplification in regions of tightly juxtaposed macrophage-target membranes where antigen-IgG-FcgR are engaged by the exclusion of phosphatases with large extracellular domains(29, 41, 44, 45). With the findings that RTX mobility increased ADCP, we sought to capture a high-resolution view of IgG-FcγR organization on the membrane within the phagocytic synapse using our previously established polarized-TIRF microscopy approach (30, 46). Here, SLBs on glass coverslips displaying IgG2a-AF647 docked to DSPE-biotin were created with fluid-phase POPC or gel-phase DPPC phospholipids. FRAP measurements confirmed that the POPC was fluid with a diffusion coefficient for the DSPE-biotin-IgG-AF647 of ∼0.93 um^2^/s (Fig. 3A, B). TIRF microscopy of DiI-labeled FLMs provided an exquisite view of the movements of the plasma membrane at large, and IgG-AF647 specifically, which is governed by its engagement with FcgRs (Supplemental Fig. 3). Consistent with our previous observations (30), interaction of the macrophage with mobile IgG resulted in the formation of focused IgG microclusters that transitioned into rapidly advancing lamellar sheets with a leading edge studded with small IgG microclusters (Fig. 3C, Video 5). A fraction of these microclusters would aggregate together and detach from the leading edge, mirroring the movements of RTX microclusters observed in confocal and LLS microscopies. On a mobile POPC bilayer, only narrow bands and patches of macrophage plasma membrane that were marked with microclusters of IgG-AF647 were observed within the ∼150 nm deep TIRF field, indicating that much of the macrophage plasma membrane was positioned far from the glass surface and was not illuminated by the TIRF field (Fig. 3C, Video 5). In contrast, IgG-AF647 on gel-phase DPPC SLBs did not move, and despite some microheterogeneities in the SLB, the macrophage membrane produced comparatively uniform DiI fluorescence in the TIRF field, indicating a tight apposition to the SLB (Fig. 3D, Video 6). Surfaces with mobile IgG-AF647 restricted IgG to the front of the advancing pseudopod. Most of the plasma membrane was far from the glass, reflecting either the distribution of tall proteins that supported it above the SLB or membrane tension created by actin tethers experiencing contractile force.

**Figure 3.**
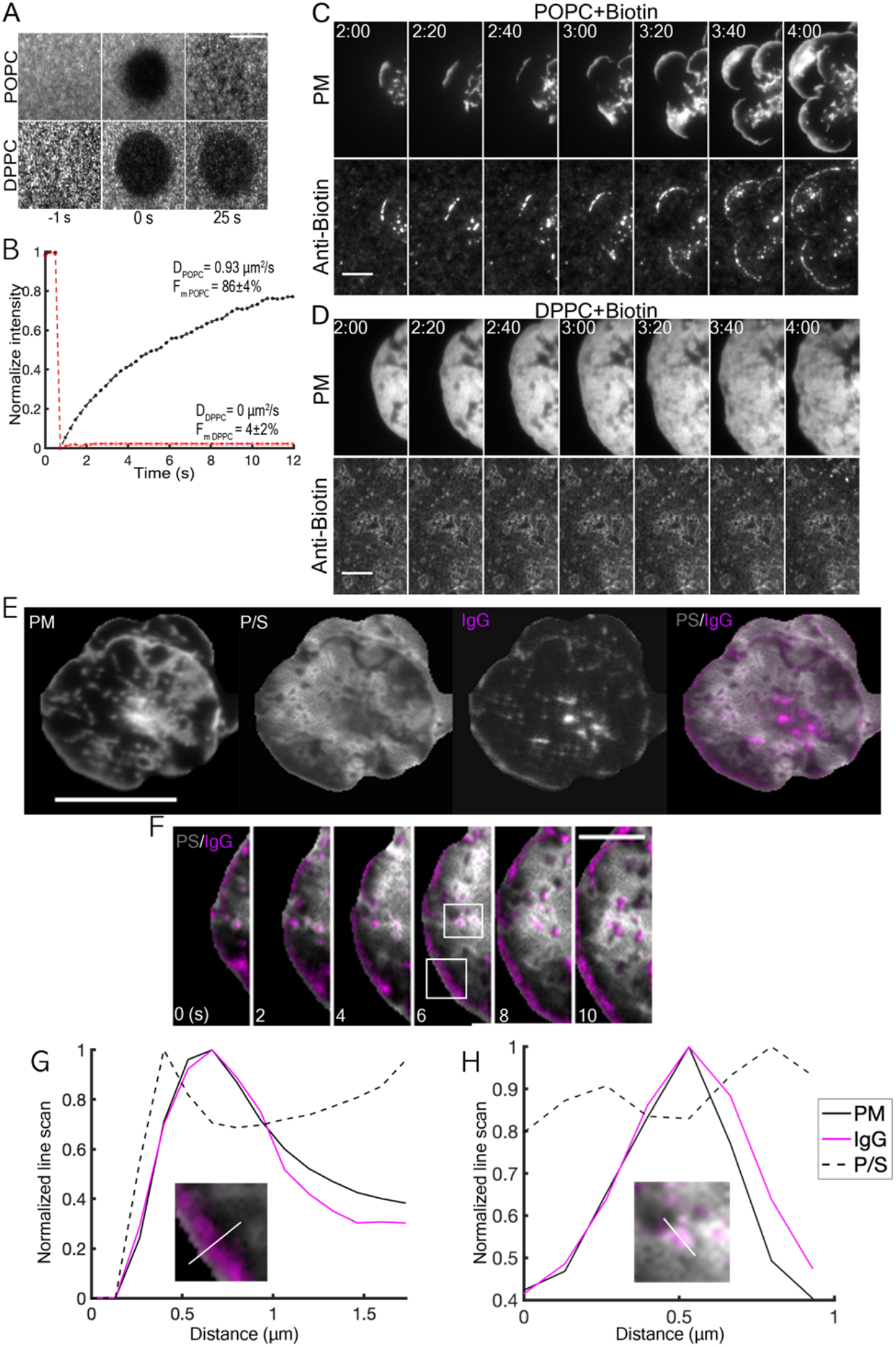
Membrane topography defines microcluster formation and patching of mobile IgG. (**A**) TIRF-FRAP of IgG-AF647 presented on mobile/fluid POPC or gel-phase/immobile DPPC SLBs. Scale bar = 5 µm. (**B**) FRAP recovery curves and fitted diffusion coefficient (D) and mobile fraction (F_m_). (**C and D**) TIRF imaging of IgG-AF647 relative to FLM plasma membrane (DiI) during engagement of IgG presented on mobile/fluid POPC (**C**) or gel-phase/immobile DPPC (**D**) SLBs (scale bar = 10 µm). (**E**) Pol-TIRF imaging of an FLM labeled with DiI relative to mobile IgG-AF647 presented on fluid POPC SLB. PM is the total DiI signal (P+2S), and membrane curvature is observed in the P/S ratio (high intensity = vertical membrane, low intensity = flat) relative to IgG-AF647 microclusters (magenta). Scale bar = 10 µm. (**F**) A zoomed region of the leading edge illustrates IgG-AF647 formation during pseudopod advancement and development of podosome-like structures detaching from the leading edge. Scale bar = 5 µm. (**G and H**) Linescan of the leading edge (**G**) indicated tight apposition of the PM to the glass where IgG-AF647 microclusters are forming in a moving band (from top boxed region in F). Linescan across a podosome-like structure (**H**), indicated maximal IgG-AF647 signal on a flat patch of PM (P/S valley), surrounded by vertical membrane (high P/S, from bottom boxed region in F).

To gain a better understanding of the membrane topography, we applied polarized-TIRF (pol-TIRF) microscopy to image changes in the topography of the plasma membrane. Previously we applied pol-TIRF to detect membrane curvature during clathrin-mediated endocytosis (46), demonstrating its excellent sensitivity to changes in membrane topography, even on objects below the diffraction limit of light. In pol-TIRF, DiI embedded in the plasma membrane orients its dipole moment with the plane of the macrophage plasma membrane. Thus, pol-TIRF parallel (s-pol, S) to the glass excites horizontal membrane and perpendicular (p-pol, P) to the coverslip excites vertical membrane. The overall nanoscale membrane topography is encoded in the P/S ratio image, in which bright regions indicate vertical plasma membrane and dim regions indicate horizontal (Fig. 3E, Video 7). Comparison of the P/S image with the distributions of plasma membrane IgG-AF647 provided a high-resolution view of the contact topography of the phagocytic synapse (Fig. 3E). The leading edge of the advancing pseudopod displayed high P/S ratio, consistent with the rapid transition in membrane topography (Fig. 3F). This narrow band of vertical membrane was immediately followed by an advancing band of newly forming IgG-AF647 microclusters (Fig 3F,G). The membrane immediately behind these microclusters maintained a low P/S ratio for variable distance behind the advancing band of IgG-AF647 microclusters (indicated by a dim P/S region) following which there was an increase in P/S and decrease in total DiI signal, signifying a rise away from the contact site (Fig. 3F and 3G). Periodic deviations of the flat membrane to vertical were observed during which intense patches of IgG-AF647 would coalesce within the flat region and be surrounded by rings of high P/S, reminiscent of podosomes (Fig. 3F and 3H). Taken together, we interpret this result to mean that the leading edge was held in tight apposition to the SLB by the IgG-AF647, followed by a band of polymerizing actin filaments that propelled the IgG-AF647 forward during expansion and maintained contact of the trailing membrane with the SLB surface. The IgG-AF647 microclusters would then disassociate from the leading edge and assemble into podosome-like structures that moved centripetally toward the center of the macrophage-contact site, where they would accumulate into larger patches as maximal spreading was reached (Supplemental Fig. 4). These movements mirrored the microclustering behavior of IgG-AF647 during engagement with WIL2-S cells. There are a number of notable differences however, that include: a population of microclusters was maintained at the leading edge, potentially mimicking the engulfment phase. Thus, translating the pol-TIRF results into the 3D context of ADCP, we conclude that the local membrane topography is defined by by small IgG-FcgR microclusters contacts propelled at the leading edge of the pseudopod and by podosome-like patches of IgG-FcgR that protrude from the macrophage plasma membrane.

### IgG-FcγR microclusters organization and pseudopod expansion require Arp2/3-mediated actin branching and localized Syk kinase activity

With the finding that pTyr and pSyk were elevated in phagocytic cups engaging mobile SLB-SBs relative to immobile SLB-SBs (Fig. 1E,F), and striking recruitment of Syk to RTX-FcgR microclusters during ADCP (Fig. 1), we speculated that these microcluster patches would serve as signal amplification hot spots for signals promoting pseudopod advancement (47). Consistent with our previous work (30), TIRF imaging demonstrated that Syk-mScarlet or AktPH-mScarlet displayed maximal intensity coincident with IgG-AF647 microclusters (Fig. 4). In contrast, Syk-mScarlet expressing FLMs on DPPC-SLBs showed near-perfectly uniform Syk-mScarlet intensity along the tightly adhered FLM plasma membrane (Fig. 4B), suggesting that the IgG density was sufficient to allow activation of FcγRs without the creation of discrete microclusters. However, the absolute fraction of Syk recruited to the membrane between cells cannot be directly compared using these methods. AktPH-mScarlet, which binds the PI3K products PI(3,4,5)P_3_ and PI(3,4)P_2_ (48) with similar affinities, showed that on fluid POPC bilayers that PI3K activity was enriched at microclusters along the leading edge and at accumulated IgG-AF647 patches (Fig. 4C), as well as diffuse accumulation on the plasma membrane. We interpret this pattern to indicate that the microclusters likely serve as PI3K activity hot spots, from which the PI(3,4,5)P_3_ and PI(3,4)P_2_ diffuse. In comparison, IgG-AF647 presented on immobile DPPC-SLBs also recruited AktPH-mScarlet to the plasma membrane more uniformly but containing some dim punctate regions (Fig. 4D). Importantly, pseudopod expansion was more extensive on mobile IgG/POPC than immobile/DPPC SLBs with FLM cells on fluid IgG expanding to cover approximately double the surface area (Fig. 4G). From these findings, we conclude that the mobile IgG allows for extensive FcγR microcluster formation and signaling via Syk and PI3K leading to more expansive pseudopod extension.

**Figure 4.**
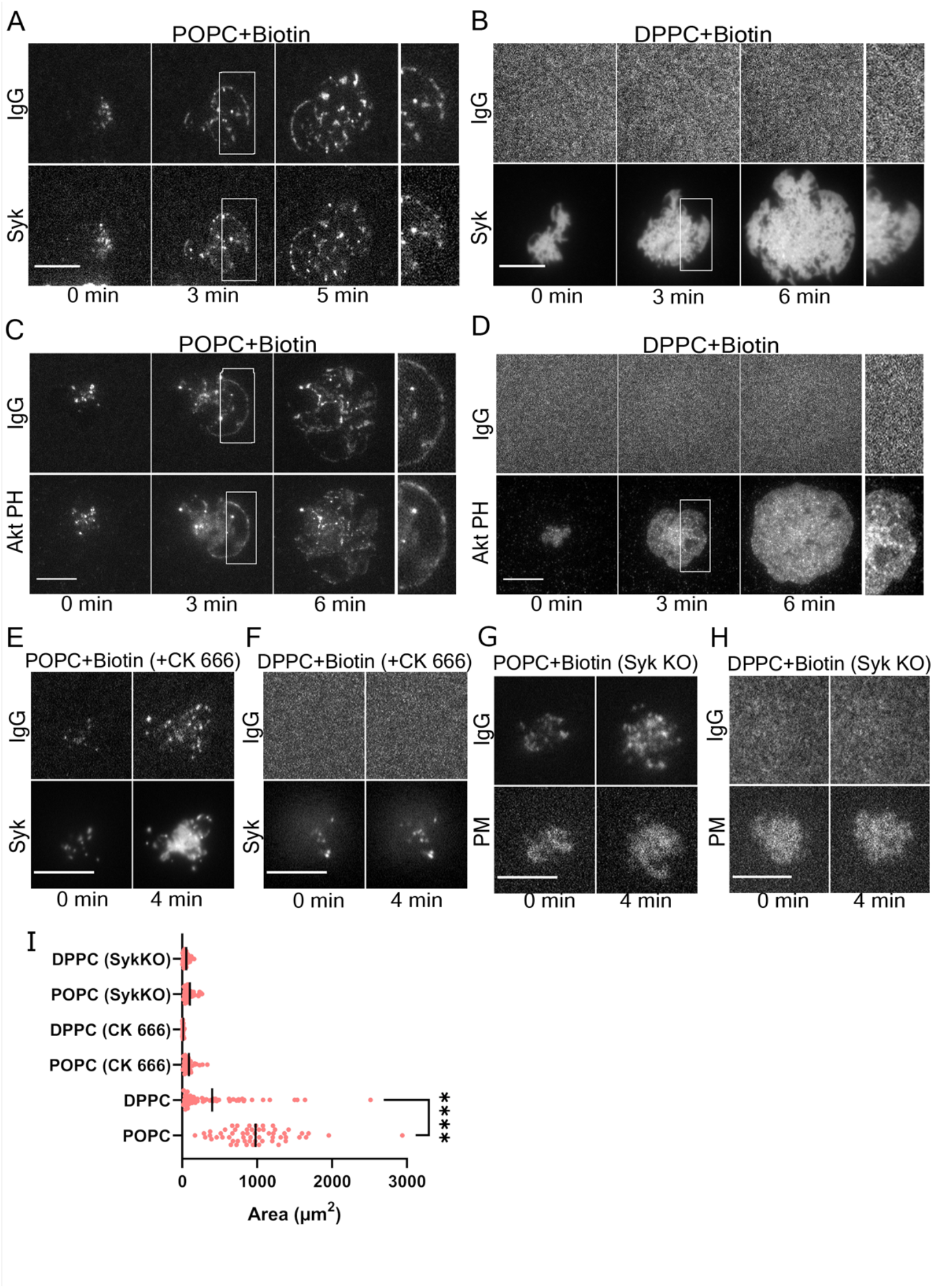
Syk and PI3K signaling originates from FcγRs IgG microclusters and is supported by an Arp2/3-Syk amplification loop. (**A-B**) TIRF imaging of FLM cells expressing Syk-mScarlet spreading on IgG-AF647 presenting mobile/fluid POPC or immobile/gel-phase DPPC SLBs. (**C-D**) AktPH-mScarlet domain expressed in FLM cells engaging IgG-AF647 presented on POPC or DPPC bilayers. (**E-F**) The localization of Syk-mScarlet with IgG-AF647 microclusters on POPC and DPPC bilayers. (**G-H**) Distribution of IgG-AF647 presented on POPC or DPPC bilayers engaged by Syk KO FLM cells expressing mNG-Mem. All scale bars are 10 µm. (**I**) Quantification of FLM cell spreading on IgG-SLB-SBs (t-test comparison, bars are mean, **** p<0.001).

The topography of the IgG-FcγR microclusters at the pseudopod leading-edge (Fig. 3), combined with the observations that Arp2/3 mediates rearrangements of the B cell receptor and T cell receptors (36, 37), suggested a similar Arp2/3 dependent mechanism was responsible. To delineate how Arp2/3 contributed to IgG-FcγR microclustering and signaling, we examined the recruitment of Syk-mScarlet and cell spreading in FLMs treated with CK-666, a potent Arp2/3 inhibitor (36, 49). CK-666 treated FLMs failed to form lamellipodia on either POPC or DPPC SLBs (Fig. 4 E and F) and both demonstrated a dramatic defect in spreading compared with untreated FLMs (Fig. 4G). Moreover, initial patches of IgG-AF647 that recruited Syk-mScarlet were observed in CK-666 treated cells on both POPC and DPPC SLBs (Fig. 4E,F); however, these structures produced few additional microclusters and completely failed to reorganize into productive lamella in the absence of Arp2/3 function. Thus, the Arp2/3 complex mediated actin branching promotes spatial organization of the IgG-FcγR spatial organization and pseudopod expansion.

Given the central role of the Syk kinase in mediating signaling from the FcγR to actin and a reciprocal relationship of actin influencing FcγRs mobility and engagement, we sought to determine Syk’s role in forming and organizing microclusters in the SLB-IgG system (50, 51). Here, we generated FLMs with gene disruptions in Syk using the CRISPR/Cas9 system and transduced them with a plasma membrane probe (PM, mNG-Mem) (34). FLM Syk^KO^ cells formed disorganized contacts with the both the POPC and DPPC IgG presenting bilayers (Fig. 4G,H and 4I), similar to Arp2/3 inhibited cells. Diffuse patches of IgG-AF647 were observed on FLM Syk^KO^ as they contacted POPC SLBs (Fig. 4G), indicating that some of the FcRs were able to bind IgG-AF647; however, no organized lamellapod or spreading was observed. Moreover, FLM Syk^KO^ cells did not spread on IgG-AF647 presented on DPPC SLBs (Fig. 4H), suggesting that FcγR is necessary to rearrange the cytoskeleton to permit engagement and additional FcγR binding. Quantification of cell spreading indicated that the diameter of macrophage spreading was twice as great on mobile IgG SLBs vs. immobile and that this response required Syk and Arp2/3 to the same degree (Fig. 4I). Together, these results suggest that Arp2/3 directs microcluster formation through actin branching which in turn enables Syk-mediated amplification of IgG-FcγR signaling. Moreover, mobile IgG facilitates microclustering of IgG-FcγRs to potentiate signaling driving pseudopod extension and reinforcement through the formation of additional IgG-FcγR binding and microclustering. Thus, similar to the rearrangement of the BCR and TCR, Arp2/3 is necessary for organizing and rearranging IgG-FcγR microclusters as well as formation of the lamellar pseudopod (36, 37).

### Failed ADCP is characterized by the trogocytic clearance of RTX at the base of the phagocytic cup during insufficient signal amplification by Syk and Arp2/3

Failed ADCP is an important problem for the efficacy of therapeutic antibodies owing to antigenic modulation and failure to kill (12, 16, 52). To gain a better understanding of the movements of IgG during failed ADCP, we analyzed confocal movies of FLMs engaging WIL2-S cells displaying confined (non-treated) or fully mobile (LatB treated) RTX. In cases where ADCP failed, RTX-AF647 accumulated at the base of the phagocytic cup where it formed patches that were internalized into a mixture of small and large vesicles for non-treated WIL2-S (Fig. 5A, Video 8) or predominantly into a larger single vesicle for LatB treated WIL2-S cells (Fig. 5B, Video 9). In comparing complete ADCP with this trogocytic failure, we observed that successful events involved a propelling of the RTX microclusters forward over the particle during the expansion of the phagocytic pseudopod (compare Figs. 5A,B with 2E and 1A,B and Videos 8,9 and Videos 1,2). We speculated that the trigger for ADCP was governed by the degree of Syk-mediated phosphorylation at these microcluster sites. Indeed, LLSM showed that Syk^KO^ FLMs, which were defective for ADCP, readily pooled RTX-AF647 at the base of phagocytic cup leading to trogocytic clearance of the RTX (Fig. 5C, Video 10). Thus, Syk activity is required to drive extension of the pseudopod and rearrangement of the microclusters within that structure. Importantly, trogocytic biting appeared to be an active process that also removed a portion of the target cell’s plasma membrane (Fig. 5D, Video 11), consistent with it causing some damage to the target cell (15). To determine if our findings for the importance of Arp2/3 translated to cellular targets, we imaged FLM cells treated with CK-666 and challenged with RTX opsonized WIL2-S cells. Indeed, these cells showed no discernable Syk-mScarlet localization to the phagocytic cup and they trogocytosed RTX similar to Syk^KO^ (Fig. 5E). This behavior was consistent across movies of 8 unique phagocytic cups. Taken together, we conclude that Syk and Arp2/3 work together to amplify signals produced by IgG-FcγRs to drive to engulfment and rearrangement of IgG-FcγRs.

**Figure 5.**
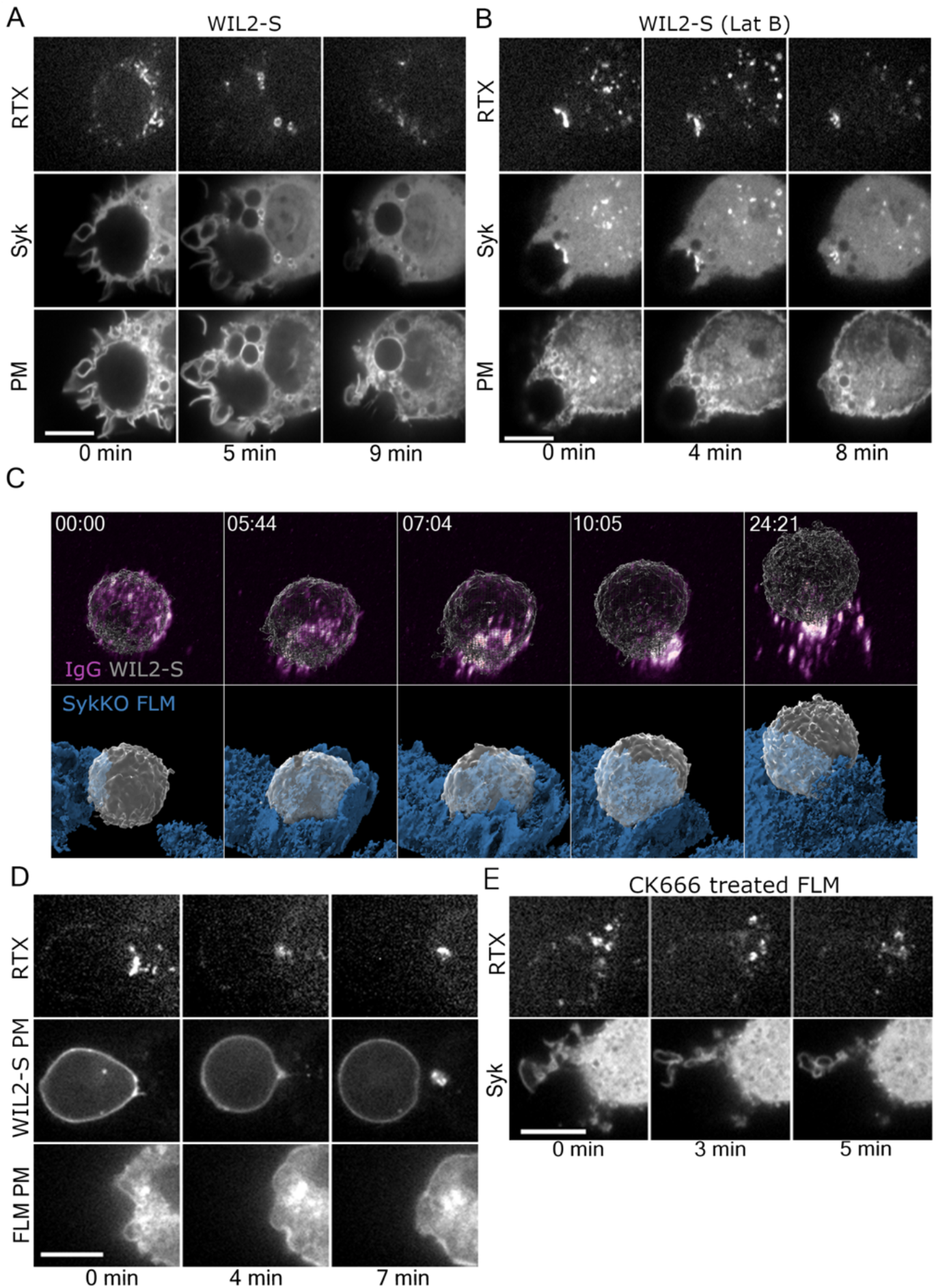
Trogocytosis is characterized by a clearance of RTX microclusters at the base of the phagocytic cup and failure to redistribute them during engulfment. (**A and B**) Confocal imaging of the movements of RTX-AF647 during trogocytosis of WIL2-S cells by FLMs without (**A**) and with LatB treatment (**B**). FLMs expressing mNG-Mem and Syk-mScarlet illustrate that, during trogocytosis, RTX recruits Syk-mScarlet to patches at the base of the cup; however, these patches are internalized into multiple large endosomes (**A**) or fewer even larger endosomes when the target is treated with LatB (**B**) (scale bar = 10 µm). (**C**) LLSM imaging of trogocytosis of a WIL2-S cell labeled with calcein violet and opsonized with RTX-CF647 by a SykKO FLM expressing mNG-Mem. Initial microclustering and formation of an RTX-CF647 ring was observed (5:44, 7:04 min); however, unlike WT FLMs, pseudopod extension fails (blue surface) and the RTX-CF647 is cleared from the WIL2-S cell and packaged into large endosomes within the FLM cell (Image dimension = 20 µm x 20 µm). (**D**) Membrane transfer from WIL2-S cell during trogocytosis. Confocal imaging of WIL2-S cells expressing mScarlet-Mem and RTX-AF647 engaging with an FLM expressing mNG-Mem. (**E**) Trogocytosis by a CK666 treated FLM showed little recruitment of Syk-mScarlet to the RTX-AF647 clusters or endosomes. All scale bars = 10 µm.

**Figure 7.**
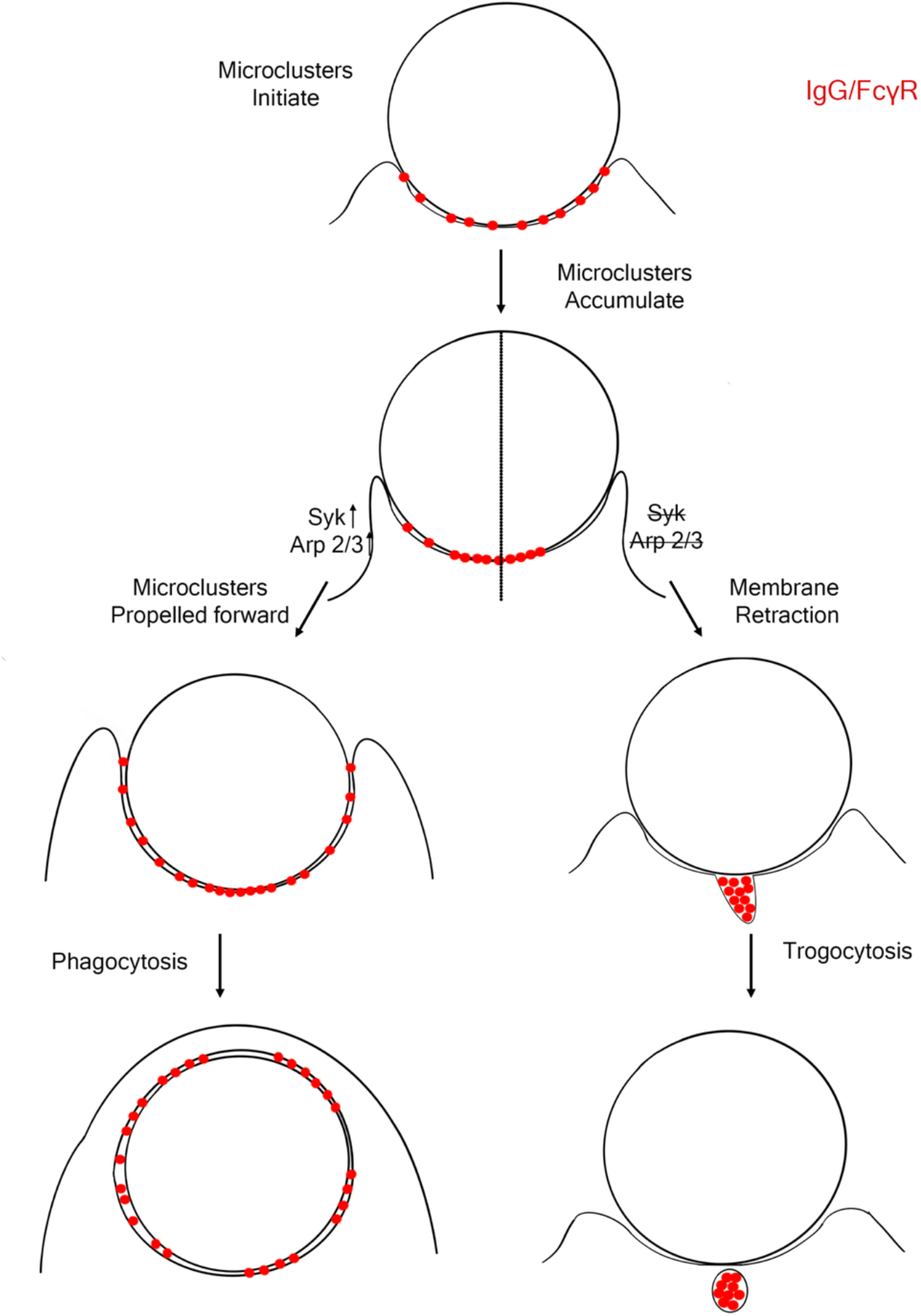
ADCP and trogocytosis are dependent on rearrangement of IgG-FcγR microclusters at the contact site. The movements of IgG-FcγR microclusters are represented by small red circles. During ADCP, IgG-FcγR microclusters are formed during contact with the target. These microclusters co-patch and begin to pool at the at the base of the phagocytic cup where the continue to recruit Syk. On the case of ADCP, sufficient Syk and Arp2/3 activity drives the microclusters forward during engulfment resulting in their nearly uniform distribution, attaching the phagosome membrane to the target. Insufficient Syk or Arp2/3 activity results in trogocytic clearance of the IgG-FcγR microclusters or patches, membrane retraction and ultimately release of the target cell.

## Discussion

The finding that IgG-FcγRs undergo substantial spatial rearrangements that potentiate signaling promoting ADCP has significant ramifications for understanding the cellular mechanisms of phagocytosis and the development of therapeutic antibodies and their adjuvants. The current paradigm submits that high densities of immobile IgG on the target cell/particle promote short-range clustering of FcγRs leading to phosphorylation by Src family members and the recruitment of Syk(53), and that this process is reinforced as the advancing pseudopod zippers along this surface (25–27, 54). While the zipper model likely holds for many rigid particles, including cell walled gram-positive bacteria and yeast, our results demonstrate that for CD20 on lymphocytes, and by extension cancer cells, or virally infected cells displaying surface mobile antigens, that the trigger model of ADCP needs to be revisited (27). The significant short- and long-range reorganization of IgG-FcγRs complexes suggests a previously underappreciated spatial regulation of FcγR signal amplification and a bifurcation in ADCP and trogocytosis with new functional roles implicated for Syk and the Arp2/3 complex (Fig. 6). Most notably, the formation of microclusters followed by pooling of RTX-FcγR toward the base of phagocytic cup is highly reminiscent of the stages of immune synapse formation in T cells (45). Unlike the TCR immune synapse which accumulates receptors into a stabilized structure; however, super-threshold signaling from the FcγRs leads to explosive pseudopod extension and target cell engulfment and the disbursement of the FcγRs over the target, tethering the phagosome membrane to it (Fig. 1C, Video 2). The formation of IgG patches by the coalescence of microclusters fits with the well-supported biochemical arguments that FcγRs must be grouped on the cell surface to allow constitutive phosphorylation by Src family kinases, including Lyn, to exceed the dephosphorylation by CD45,

CD148 and others to generate the phosphorylated-ITAMs that allow docking of Syk (29, 45, 55). Thus, grouping into these patches amplifies FcγRs signaling and lowers the threshold needed to ‘trigger’ the engulfment of the target cell. This is consistent with our observations of enhanced ADCP by disruption of the target cell’s cytoskeleton thereby increasing RTX mobility, lowering the density of IgG required for ADCP and increasing the number of target cells engulfed (Fig 2B). A direct implication of this finding is that target cells with dense or active actin networks that restrict the mobility of surface antigens are likely to be increasingly resistant to ADCP, requiring higher densities of therapeutic antibody. In corollary, treatments that depolymerize or somehow disrupt actin corralling of surface antigens on target cells may act as adjuvants for antibody therapies acting through ADCP.

While the exact cellular mechanisms that promote microclustering of the IgG-FcγR complexes are still unclear, this study implicates a positive feedback loop involving Arp2/3 and Syk. Arp2/3 mediated actin branching, similar to that observed in spatial patterning of the BCR (36), likely plays a critical role in enhancing the FcγR signaling by driving assembly of microclusters near the leading edge. While we have not characterized the actin defects resulting from Arp2/3 in this study, inhibition of Arp2/3 has a dramatic effect on the formation and reorganization of IgG-FcγR microclusters to SLBs and led to trogocytosis rather than ADCP. We interpret these findings to mean that actin branching coupled with retrograde flow is important at the earliest stages of FcγR microcluster formation and likely contributes to the proximal grouping of these receptors necessary for Src family mediated phosphorylation and Syk recruitment. Likewise, the transition of IgG-FcγRs into larger patches may involve other actin remodeling factors including the Arp2/3 complex for the formation of more podosome like actin structures (56). Based on our LLSM imaging, we would also conclude that sufficient FcγR signal amplification to trigger ADCP leads to disbursement of these microclusters during engulfment, suggesting a potential second role for actin polymerization in their reorganization. These findings predict that changes in the macrophage transcriptional program that potentiate Syk signaling or Arp2/3 activity potentially through enhanced activation of the Arp2/3 nucleators WAVE2 and WASP, should also be strong predictors of ADCP.

## Materials and Methods

### Cell culture

Fetal liver-derived macrophages (FLMs) were prepared from E18 mouse fetuses from B6J.129(Cg)-Igs2tm1.1(CAG-cas9*)Mmw/J mice (The Jackson Laboratory, Stock No. 028239, Bar Harbor, ME) with South Dakota State University Institutional Animal Use and Care Committee approval. Liver tissue was mechanically dissociated using sterile fine-pointed forceps, and a single cell suspension was created by passing the tissue through a 1mL pipette tip. Bone marrow-derived macrophages (BMDMs) were prepared using single cell suspensions of bone marrow obtained from the long bones of B6J.129(Cg)-Gt(ROSA)26Sortm1.1(CAG-cas9*,-EGFP)Fezh/J mice (The Jackson Laboratory, Stock No. 026179, Bar Harbor, ME). BMDM and FLMs were cultured in bone marrow media containing DMEM (Corning) supplemented with 30% L cell supernatant, 20% Heat-inactivated FBS (R&D Systems) and 1% pen/strep (ThermoFisher scientific) and incubated at 37°C and 5% CO_2_. Prior to LLSM experiments FLMs were cultured in DMEM/F12 (Gibco) supplemented with 20% heat-inactivated FBS (R&D Systems), 1% penicillin-streptomycin (Gibco), 50 ng/ml CSF-1 (BioLegend), and 0.5 µg/mL plasmocin (Invivogen) at 37° with 7.5% CO_2_. WIL2-S cells (CRL-8885) were obtained from ATCC and cultured in RPMI-1640 medium (ATCC 30-2001) containing 10% FBS and 1% pen/strep (ThermoFisher scientific) at 37°C with 5% CO_2_.

### Plasmids, lentiviral transduction and CRISPR gene disruption

Lentiviral expression plasmids, pLJM1-AktPH-mScarlet, pLJM1-Syk-mScarlet, and pLJM1-Lck(mem)-mNeonGreen were created by gene synthesis (GenScript, Piscataway, NJ). pCMV-VSV-G was a gift from Bob Weinberg (Addgene plasmid #8454; http://n2t.net/addgene:8454; RRID:Addgene_8454) (57). psPAX2 was a gift from Didier Trono (Addgene plasmid #12260; http://n2t.net/addgene:12260; RRID:Addgene_12260). pLJM1-EGFP was a gift from David Sabatini (Addgene plasmid #19319 ; http://n2t.net/addgene:19319 ; RRID:Addgene_19319)(58). Lentiviral vectors were produced in 293T cells by transfection with pLJM1-Syk-mScarlet-I and pLJM1-AktPH-mScarlet-I in combination with helper plasmids psPAX2 and pCMV-VSV-G, using linear 25 kDa polyethylenimine (PEI). Lentiviral supernatant was harvested after 48 hours, centrifuged, and added to FLM cells cultured in bone marrow media supplemented with 10 µg/ml of cyclosporine A. Virally transduced cells were selected with Blasticidin (10 µg/mL) and/or Puromycin (5 µg/mL) for 48 hours post-transduction. Macrophages were split into respective well formats for experiments. Once BMMs or FLMs reached appropriate seeding density, they were used for phagocytosis assays, frustrated phagocytosis, and live imaging.

### Antibody labeling with NHS ester fluorophores

Anti-human CD20 murine IgG2a (hcd20-mab10, InvivoGen) and mouse anti-Biotin mIgG2a [3E6] (ab36406, Abcam) were fluorescently labeled using Alexa Fluor 647 NHS ester (Thermo Fisher Scientific) or CF 647 NHS ester (Biotium). Prior to labeling, antibodies were buffer exchanged in Zeba Spin desalting columns (ThermoFisher Scientific) with three washes of PBS, pH 7.4 without Ca^2+^ and Mg^2+^. Exchanged antibodies were mixed with NHS-fluorophore at molar ratio of 1:10 IgG:fluorophore in 0.1M sodium bicarbonate solution (ThemoFisher Scientific) with pH of 8.1-8.3. The mixture was incubated for 1 hour at room temperature. After 1 hour incubation, excess fluorophores in the solution were removed using Zeba Spin columns (ThemoFisher Scientific).

### Supported lipid bilayers

Bilayers were formed by spontaneous fusion of lipid vesicles. Small unilamellar vesicles (SUVs) were prepared by mixing DSPE-PEG(2000)-Biotin (880129 Avanti Polar Lipids) and DPPC (850355, Avanti Polar Lipids) or POPC (850457, Avanti Polar Lipids) at a molar ratio of 1:1000 with total lipid concentration of 500 µM (380 µg/mL) in chloroform. Lipids were then roto-vacuum dried for up to 60 min in a glass test tube. The lipid film was resuspended using 1 mL PBS, pH 7.4 without Ca^2+^ and Mg^2+^. The glass tube was then sonicated for 5 minutes using a bath sonicator (JSP US40) followed by extrusion (610000, Avanti Polar Lipids) through a 100 nm filter at least 13 times (Whatman Nucleopore Track-Etch 100 nm membrane). POPC liposomes, T_m_ of -2°C, were extruded and sonicated at room temperature, whereas DPPC liposomes with T_m_ of 41°C, were extruded and sonicated at 60°C.

The 500 µM stocks of SUVs were diluted 1:6 in 2mM Mg^2+^ PBS. POPC supported lipid bilayers (SLBs) were formed by pipetting the diluted lipids onto Hellmanex III (MilliporeSigma) cleaned 96-well (Dot Scientific, MGB096-1-2-LG-L) plates and incubated at 37°C for 30 minutes. DPPC bilayers were formed by pipetting the diluted lipids onto 25 mm Hellmanex III (MilliporeSigma) cleaned coverslip mounted in a pre-warmed Attofluor chamber (Thermofisher Scientific) and incubated at 70°C for 30 minutes on a slide-warmer (Premier XH-2001). Bilayers were also formed on silica beads with 5 µm diameter (Bangs Laboratories). Silica beads (SB) were washed by centrifugation twice with acetone, twice with ethanol and twice with deionized water. POPC and DPPC SUVs were then incubated at 37°C or 70°C for 30 minutes with the cleaned silica beads. Following incubation, excess liposomes were removed by pipetting in Hank’s Balanced Salt Solution (HBSS, Corning™ 21023CV) containing FBS in 1:1000 ratio (HBSS+0.1% FBS). SLBs and SLB-SBs were incubated with anti-Biotin IgG (3E6) (Abcam ab36406) conjugated with Alexa Fluor 647 NHS ester (Thermo Fisher scientific) SLB at 37 °C for 30 min. SLBs were rinsed without exposure to air to remove excess IgG using HBSS.

For imaging FLM cells on SLBs, SLBs were made directly on Hallmanex III (MilliporeSigma) cleaned 25 mm coverslips (Number 1.5, Thermo Fisher) or onto 96-well glass bottom plates (Dot Scientific, MGB096-1-2-LG-L) for high content experiments. The 25 mm coverslips were imaged in AttoFluor chambers (Thermo Fisher).

### Immunostaining of pTyr and pSyk

BMDMs were plated into 96 well glass bottom plates and exposed to silica beads bearing DSPE-PEG(2000)-Biotin at 1:1000 in POPC or DPPC bilayers and opsonized using anti-Biotin IgG2a for 12 minutes. Cells were then rinsed with ice cold PBS and fixed in 4% PFA at room temp for 30 minutes, followed by blocking with PBS containing 5% FBS for 10 min and incubation with a 1:50 dilution of anti-pSyk-AF647 (C87C1) or anti-pTyr-AF647 (P-Tyr-100) and anti-CD45-AF488 (D3F8Q, Cell Signaling Technology) overnight at 4°C with gentle rocking. Images were captured using an ImageXpress Micro XLS Widefield High-Content Analysis System (Molecular Devices) using a 20x 0.70 N.A. objective lens.

### Manipulation of actin and CD20 mobility in WIL2-S cells

Paraformaldehyde (PFA, T353-500, ThermoFisher scientific), Latrunculin B (Lat B, 428020, MilliporeSigma) and Jasplakinolide (Jas, 420107, MilliporeSigma)/Blebbistatin (Bleb, B0560, Sigma-Aldrich) were used for manipulating CD20 mobility. 4% PFA was prepared using powder form PFA which was diluted to prepare 2% PFA in PBS, pH 7.4 without Ca^2+^ and Mg^2+^and stored at -20°C. WIL2-S cells were treated using 2% PFA for 10 minutes at 4°C. Lat B was resuspended in DMSO at a concentration of 25 mg/mL (62.5 mM) and stored at - 20°C. 12.5 mM Lat B solution was prepared from the stock solution and used to treat WIL2-S cells for 30 minutes at 4°C. Jas and Bleb were stored in DMSO with concentrations of 0.71 mg/mL (1mM) and 5 mg/mL (17mM), respectively, at - 20°C. Cells were incubated with 75 µM Bleb for 10 minutes, followed by 1µM Jas for 5 minutes. After drug treatment, WIL2-S cells were thoroughly washed by centrifugation 3 times and incubated on ice.

FLMs were treated with the Arp2/3 inhibitor CK-666 (MilliporeSigma) for live imaging. Commercially purchased CK-666 was stored in DMSO with a concentration of 25 mg/mL (85mM) at 4°C. FLMs were treated with 1.2 µM CK-666 in HBSS for 1 hour at 4°C before live imaging.

### TIRF imaging

FLMs were dropped onto prepared SLBs. TIRF-based imaging was conducted using an inverted microscope built around a Till iMic (Till Photonics, Germany) equipped with a 60 × 1.49 N.A. oil immersion objective lens. The entire microscope setup and centering of the back focal plane was previously described in (46). Cells were labeled by rinsing with 1μg/mL DiI in 2.5% DMSO/PBS. Excitation for TIRF was provided by a 633nm laser for AF647 anti-biotin mIgG2a, a 561 nm laser for DiI, Syk-mScarlet, and AktPH-mScarlet, and a 488 nm laser for Mem-mNG.

TIRF 360 was used to create uniform TIRF illumination for fluorescence recovery after photobleaching (FRAP) by steering the laser at the back-focal plane as described (59). For spreading area, wild type, Syk KO and CK-666 treated FLMs were dropped on both mobile/fluid and immobile/gel-phase SLBs. FLMs expressing Mem-mNG were imaged 10 minutes after dropping, and spreading areas were measured by manually thresholding Mem-mNG fluorescence using ImageJ/Fiji (60).

### Confocal imaging

FLMs plated on ethanol flamed 25 mm coverslips were challenged with biotin labeled and IgG2a opsonized WIL2-S cells as targets. Imaging was performed on a TIL photonics Andromeda Spinning Disk Confocal (TILL Photonics, Munich, Germany), using a 60x 1.4 N.A. oil immersion objective lens. Excitation for confocal imaging was provided by a 640 nm laser for AF647 anti-CD20 mIgG2a, a 561 nm laser for Syk-mScarlet and Mem-mScarlet, and a 515 nm laser for mNG-Mem.

### High content imaging

For high content experiments, BMDMs were plated on 96-well plates for at least 2 hours (Dot Scientific, MGB096-1-2-LG-L) before imaging, WIL2-S cells were opsonized with anti-hCD20 mIgG2a (InvivoGen, RTX) and treated with different drugs including PFA, Lat B and Jas/Bleb. Fluorescent detection of internalized targets was achieved by labeling WIL2-S cells with pHrodo™ SE Red dye (ThermoFisher scientific) in PBS, PH 7.4 without Ca^2+^ and Mg^2+^. Media was replaced with HBSS prior to imaging by pipetting. High content fluorescent imaging was conducted using an ImageXpress Micro XLS Widefield High-Content Analysis System (Molecular Devices, Sunnyvale, CA) equipped with a 20 × 0.70 N.A. objective lens. For each well, 4 fields of view were imaged for pHrodoRed (ex. 562/40, em. 624/40), GFP (ex. 482/35, em. 536/40) and HCS NuclearMask Blue (ex. 377/50, em. 447/60). Images were analyzed using a custom pipeline in CellProfiler (61) to obtain the number of engulfed WIL2-S cells per macrophage to quantify the phagocytic index (PI).

The density of IgG on cells was measured using Alexa Fluor 633 Goat anti-Mouse IgG2a Cross-Adsorbed Secondary Antibody (ThermoFisher scientific). The secondary antibody was diluted 1:1000 in HBSS+0.1% FBS. WIL2-S cells were opsonized with dilutions of anti-human CD20 – murine IgG2a (hcd20-mab10, InvivoGen) for 30 min followed by incubation with the diluted secondary antibody for 30 minutes and washed 3 times with HBSS. FACS analysis was performed on a BD Accuri™ C6 Flow Cytometer and analyzed using BD FlowJo™.

### Cell preparation for LLSM ADCP

2-3×10^5^ FLMs expressing Syk-mScarlet and Mem-mNG were plated overnight in a 6 well dish on absolute ethanol flame-cleaned 5 mm diameter #1 thickness coverslips (Warner Instruments). 1.5×10^6^ WIL2-S cells were incubated with 20µM calcein violet AM (BioLegend) and 5 µg/mL of Rituximab-CF647 for 30 min at 4°C and washed at least 3 times in HBSS with 1%FBS. The field of view was chosen to contain a single FLM and 3×10^5^ labeled WIL2-S cells were dropped on top of the coverslip and imaged immediately after to capture the initial contact.

### Lattice light sheet microscopy

Volumetric image stacks were generated using dithered square virtual lattices (Outer NA 0.55, Inner NA 0.50, approximately 30 µm long), stage scanning of 151 slices with 0.5 µm step sizes, resulting in 254 nm deskewed z-steps, a region of interest of ∼76µm x 76µm x 42µm, and 12 ms exposures resulting in 10-12 s intervals. The laser illumination was optimized to minimize photobleaching during the full acquisition, resulting in 0.58µW (405nm) for Calcein violet, 15 µW (488nm) for Mem-mNG, 22 µW (560nm) for Syk-mScarlet, and 17µW (640nm) for Rtx-CF647 as measured at the rear aperture of the excitation objective.

### Post processing and visualization

The raw volumes acquired from the LLSM were processed using a standard pipeline(62) including deskewing, deconvolving, and rotating to coverslip coordinates in LLSpy (63). We applied a fixed background subtraction based on an average dark current image, 10 iterations of Lucy-Richardson deconvolution with experimentally measured point spread functions for each excitation followed by rotation to coverslip coordinates and cropping to the region of interest for visualization. The fully processed data was opened as a volume map series in UCSF ChimeraX and utilized isosurface, mesh, and 3D volumetric renderings to exam the data.

## Visualization

The data visualization and analyses were performed using UCSF ChimeraX, developed by the Resource for Biocomputing, Visualization, and Informatics at the University of California, San Francisco, with support from National Institutes of Health R01-GM129325 and the Office of Cyber Infrastructure and Computational Biology, National Institute of Allergy and Infectious Diseases. LLSM: The Lattice Light Sheet Microscope referenced in this research was developed under license from Howard Hughes Medical Institute, Janelia Farms Research Campus (“Bessel Beam” patent applications 13/160,492 and 13/844,405).

## Movie Legends

**Video 1. Confocal imaging RTX microclustering and Syk-mScarlet recruitment**. An FLM expressing Syk-mScarlet and mNG-Mem during ADCP of an RTX-AF647 opsonized WIL2-S cell. Video duration is 7 minutes (scale bar = 10 µm).

**Video 2. LLSM movie of IgG and Syk microclustering during ADCP of a WIL2-S cell**. An FLM expressing Syk-Scarlet and mNG-Mem was challenged by WIL2-S cell labeled with calcein violet and RTX-AF647. Video duration is 11 minutes (image dimension = 20 x 20 x 20 µm).

**Video 3. LLSM movie of IgG and Syk microclustering on Lat B treated WIL2-S cell**. An FLM expressing Syk-mScarlet and mNG-Mem was challenged by WIL2-S cell labeled with calcein violet and RTX-AF647. Video duration is 11 minutes (image dimension = 21 x 21 x 21 µm).

**Video 4. LLSM movie showing lack of IgG and Syk microclustering and colocalization on PFA treated WIL2-S cell**. An FLM expressing Syk-mScarlet and mNG-Mem was challenged by WIL2-S cells labeled with calcein violet and RTX-AF647. Video duration is 10 minutes (image dimension = 25 x 25 x 25 µm).

**Video 5. TIRF imaging of IgG microclustering at the leading edge**. DiI labeled FLM on a mobile POPC SLB displaying IgG-AF647 antibody. Duration is 2 min. (scale bar = 10 µm).

**Video 6. TIRF movie of lack of IgG microclustering at the leading edge**. DiI labeled FLM on an immobile DPPC SLB displaying RTX-AF647. Duration is 2 min. (scale bar = 10 µm).

**Video 7. Pol-TIRF imaging showing membrane topography around IgG-FcγR microclusters**. DiI labeled FLM on a mobile POPC SLB displaying RTX-AF647). Vertical membrane is bright in the P/S ratio. Video duration is 2 minutes (scale bar = 10 µm).

**Video 8. Confocal movie showing trogocytosis of RTX from WIL2-S cell**. An FLM expressing Syk-mScarlet and mNG-Mem was challenged with WIL2-S cells opsonized with RTX-AF647. Video duration is 5 minutes (scale bar = 10 µm).

**Video 9. Confocal movie of trogocytosis of RTX from a Lat B treated WIL2-S cell**. An FLM expressing Syk-mScarlet and mNG-Mem with a Lat B treated WIL2-S cells opsonized with RTX-AF647. Video duration is 5 minutes (scale bar = 10 µm).

**Video 10. LLSM movie of trogocytosis**. A Syk^KO^ FLM expressing mNG-Mem was challenged by WIL2-S cells labeled with calcein violet (used to define the mesh and grey surfaces) and RTX-AF647. Video duration is 25 minutes (Image dimension = 20 µm x 20 µm).

**Video 11. Confocal movie of trogocytosis of RTX and target cell membrane**. An FLM expressing mNG-Mem was challenged by WIL2-S cells labeled with RTX-AF647 and expressing mScarlet-Mem. Video duration is 7 minutes (scale bar = 10 µm).

## References

1. M. E. Ackerman, et al., Enhanced phagocytic activity of HIV-specific antibodies correlates with natural production of immunoglobulins with skewed affinity for FcγR2a and FcγR2b. Journal of Virology 87, 5468–5476 (2013).

2. M. J. E. Marshall, R. J. Stopforth, M. S. Cragg, Therapeutic Antibodies: What Have We Learnt from Targeting CD20 and Where Are We Going? Frontiers in Immunology 8, 1327–22 (2017).

3. M. Sips, et al., Fc receptor-mediated phagocytosis in tissues as a potent mechanism for preventive and therapeutic HIV vaccine strategies. Mucosal Immunology 9, 1584–1595 (2016).

4. S. Gordan, M. Biburger, F. Nimmerjahn, bIgG time for large eaters: monocytes and macrophages as effector and target cells of antibody-mediated immune activation and repression. Immunological reviews 268, 52–65 (2015).

5. W. He, et al., Alveolar macrophages are critical for broadly-reactive antibody-mediated protection against influenza A virus in mice. Nature Communications, 1–13 (2017).

6. C. E. Mullarkey, et al., Broadly Neutralizing Hemagglutinin Stalk-Specific Antibodies Induce Potent Phagocytosis of Immune Complexes by Neutrophils in an Fc-Dependent Manner. mBio 7, 1–12 (2016).

7. C. Alvey, D. E. Discher, Engineering macrophages to eat cancer: from “marker of self” CD47 and phagocytosis to differentiation. Journal of Leukocyte Biology 102, 31–40 (2017).

8. L. N. Dahal, et al., STING Activation Reverses Lymphoma-Mediated Resistance to Antibody Immunotherapy. Cancer Research (2017) https://doi.org/10.1158/0008-5472.can-16-2784.

9. D. Kao, et al., A Monosaccharide Residue Is Sufficient to Maintain Mouse and Human IgG Subclass Activity and Directs IgG Effector Functions to Cellular Fc Receptors. CellReports 13, 2376–2385 (2015).

10. N. Gul, M. van Egmond, Antibody-Dependent Phagocytosis of Tumor Cells by Macrophages: A Potent Effector Mechanism of Monoclonal Antibody Therapy of Cancer. Cancer Research 75, 5008–5013 (2015).

11. K. Weiskopf, I. L. Weissman, Macrophages are critical effectors of antibody therapies for cancer. mAbs 7, 303–310 (2015).

12. A. K. Church, et al., Anti-CD20 monoclonal antibody-dependent phagocytosis of chronic lymphocytic leukaemia cells by autologous macrophages. Clinical and experimental immunology 183, 90–101 (2016).

13. A. T. Vaughan, et al., Activatory and Inhibitory Fcγ Receptors Augment Rituximab-mediated Internalization of CD20 Independent of Signaling via the Cytoplasmic Domain. The Journal of biological chemistry 290, 5424–5437 (2015).

14. H. L. Matlung, et al., Neutrophils Kill Antibody-Opsonized Cancer Cells by Trogoptosis. CellReports 23, 3946-3959.e6 (2018).

15. R. Velmurugan, D. K. Challa, S. Ram, R. J. Ober, E. S. Ward, Macrophage-Mediated Trogocytosis Leads to Death of Antibody-Opsonized Tumor Cells. Molecular Cancer Therapeutics 15, 1879–1889 (2016).

16. S. A. Beers, et al., Antigenic modulation limits the efficacy of anti-CD20 antibodies: implications for antibody selection. Blood 115, 5191–5201 (2010).

17. R. P. Taylor, M. A. Lindorfer, Fcγ-receptor–mediated trogocytosis impacts mAb-based therapies: historical precedence and recent developments. Blood (2015) https://doi.org/10.1182/blood-2014.

18. A. A. Barkal, et al., Engagement of MHC class I by the inhibitory receptor LILRB1 suppresses macrophages and is a target of cancer immunotherapy. Nature Immunology 19, 76–84 (2017).

19. S. R. Gordon, et al., PD-1 expression by tumour-associated macrophages inhibits phagocytosis and tumour immunity. Nature Publishing Group 545, 495–499 (2017).

20. E. C. Piccione, et al., A bispecific antibody targeting CD47 and CD20 selectively binds and eliminates dual antigen expressing lymphoma cells. mAbs 7, 946–956 (2015).

21. M. P. Chao, et al., Anti-CD47 antibody synergizes with rituximab to promote phagocytosis and eradicate non-Hodgkin lymphoma. Cell 142, 699–713 (2010).

22. H. Wang, et al., CD47/SIRPα blocking peptide identification and synergistic effect with irradiation for cancer immunotherapy. J Immunother Cancer 8, e000905 (2020).

23. S. B. Willingham, J. P. Volkmer, The CD47-signal regulatory protein alpha (SIRPa) interaction is a therapeutic target for human solid tumors in Proceedings of the …., (2012) https://doi.org/10.1073/pnas.1121623109/-/dcsupplemental.

24. M. A. Morrissey, et al., Chimeric antigen receptors that trigger phagocytosis. eLife 7, 31 (2018).

25. F. M. Griffin, J. A. Griffin, S. C. Silverstein, Studies on the mechanism of phagocytosis. II. The interaction of macrophages with anti-immunoglobulin IgG-coated bone marrow-derived lymphocytes. The Journal of Experimental Medicine 144, 788–809 (1976).

26. F. M. Griffin, J. A. Griffin, J. E. Leider, S. C. Silverstein, Studies on the mechanism of phagocytosis. I. Requirements for circumferential attachment of particle-bound ligands to specific receptors on the macrophage plasma membrane. The Journal of Experimental Medicine 142, 1263–1282 (1975).

27. J. A. Swanson, A. D. Hoppe, The coordination of signaling during Fc receptor-mediated phagocytosis. J Leukocyte Biol 76, 1093–1103 (2004).

28. T. Pham, P. Mero, J. W. Booth, Dynamics of macrophage trogocytosis of rituximab-coated B cells. PLoS ONE 6, e14498 (2011).

29. M. H. Bakalar, et al., Size-Dependent Segregation Controls Macrophage Phagocytosis of Antibody-Opsonized Targets. Cell 174, 131-142.e13 (2018).

30. J. Lin, et al., TIRF imaging of Fc gamma receptor microclusters dynamics and signaling on macrophages during frustrated phagocytosis. BMC Immunology, 1–9 (2016).

31. K. Matlawska-Wasowska, et al., Macrophage and NK-mediated killing of precursor-B acute lymphoblastic leukemia cells targeted with a-fucosylated anti-CD19 humanized antibodies. Leukemia 27, 1263–1274 (2013).

32. E. O. Saphire, et al., Systematic Analysis of Monoclonal Antibodies against Ebola Virus GP Defines Features that Contribute to Protection. Cell 174, 938-952.e13 (2018).

33. B. M. Gunn, et al., A Role for Fc Function in Therapeutic Monoclonal Antibody-Mediated Protection against Ebola Virus. Cell host & microbe 24, 221-233.e5 (2018).

34. E. M. Bailey, et al., Engineered IgG1-Fc Molecules Define Valency Control of Cell Surface Fcγ Receptor Inhibition and Activation in Endosomes. Front Immunol 11, 617767 (2021).

35. B.-C. Chen, et al., Lattice light-sheet microscopy: imaging molecules to embryos at high spatiotemporal resolution. Science 346, 1257998 (2014).

36. M. Bolger-Munro, et al., Arp2/3 complex-driven spatial patterning of the BCR enhances immune synapse formation, BCR signaling and B cell activation. eLife 8, 40 (2019).

37. S. Kumari, et al., Actin foci facilitate activation of the phospholipase C-γ in primary T lymphocytes via the WASP pathway. Elife 4, e04953 (2015).

38. P. Beemiller, J. Jacobelli, M. F. Krummel, Integration of the movement of signaling microclusters with cellular motility in immunological synapses. Nat Immunol 13, 787–795 (2012).

39. D. Rudnicka, et al., Rituximab causes a polarization of B cells that augments its therapeutic function in NK-cell– mediated antibody-dependent cellular cytotoxicity. Blood 121, 4694–4702 (2013).

40. J. Lou, S. T. Low-Nam, J. G. Kerkvliet, A. D. Hoppe, Delivery of CSF-1R to the lumen of macropinosomes promotes its destruction in macrophages. Journal of Cell Science 127, 5228–5239 (2014).

41. S. A. Freeman, S. Grinstein, Phagocytosis: receptors, signal integration, and the cytoskeleton. Immunological reviews 262, 193–215 (2014).

42. S. A. Freeman, et al., Transmembrane Pickets Connect Cyto-and Pericellular Skeletons Forming Barriers to Receptor Engagement. Cell 172, 305-317.e10 (2018).

43. A. Kusumi, K. G. N. Suzuki, R. S. Kasai, K. Ritchie, T. K. Fujiwara, Hierarchical mesoscale domain organization of the plasma membrane. Trends in Biochemical Sciences 36, 604–615 (2011).

44. S. A. Freeman, et al., Integrins Form an Expanding Diffusional Barrier that Coordinates Phagocytosis. Cell 164, 128– 140 (2016).

45. F. Niedergang, V. D. Bartolo, A. Alcover, Comparative Anatomy of Phagocytic and Immunological Synapses. Frontiers in Immunology 7, 118–9 (2016).

46. B. L. Scott, et al., Membrane bending occurs at all stages of clathrin-coat assembly and defines endocytic dynamics. Nature Communications 9, 419 (2018).

47. Y. Zhang, A. D. Hoppe, J. A. Swanson, Coordination of Fc receptor signaling regulates cellular commitment to phagocytosis. Proceedings of the National Academy of Sciences 107, 19332–19337 (2010).

48. M. Frech, et al., High Affinity Binding of Inositol Phosphates and Phosphoinositides to the Pleckstrin Homology Domain of RAC/Protein Kinase B and Their Influence on Kinase Activity*. J Biol Chem 272, 8474–8481 (1997).

49. B. Hetrick, M. S. Han, L. A. Helgeson, B. J. Nolen, Small Molecules CK-666 and CK-869 Inhibit Actin-Related Protein 2/3 Complex by Blocking an Activating Conformational Change. Chem Biol 20, 701–712 (2013).

50. R. S. Flannagan, R. E. Harrison, C. M. Yip, K. Jaqaman, S. Grinstein, Dynamic macrophage “probing” is required for the efficient capture of phagocytic targets. The Journal of Cell Biology 191, 1205–1218 (2010).

51. V. Jaumouillé, et al., Actin Cytoskeleton Reorganization by Syk Regulates Fcγ Receptor Responsiveness by Increasing Its Lateral Mobility and Clustering. Developmental Cell 29, 534–546 (2014).

52. L. N. Dahal, et al., Shaving Is an Epiphenomenon of Type I and II Anti-CD20-Mediated Phagocytosis, whereas Antigenic Modulation Limits Type I Monoclonal Antibody Efficacy. Journal of immunology (Baltimore, Md. : 1950) 201, 1211–1221 (2018).

53. A. M. Duchemin, L. K. Ernst, C. L. Anderson, Clustering of the high affinity Fc receptor for immunoglobulin G (Fc gamma RI) results in phosphorylation of its associated gamma-chain. J Biological Chem 269, 12111–7 (1994).

54. J. A. Swanson, Shaping cups into phagosomes and macropinosomes. Nature Reviews Molecular Cell Biology 9, 639– 649 (2008).

55. F. Niedergang, S. Grinstein, ScienceDirect How to build a phagosome: new concepts for an old process. Current Opinion in Cell Biology 50, 57–63 (2018).

56. S. Linder, Podosomes at a glance. Journal of Cell Science 118, 2079–2082 (2005).

57. S. A. Stewart, et al., Lentivirus-delivered stable gene silencing by RNAi in primary cells. Rna 9, 493–501 (2003).

58. Y. Sancak, et al., The Rag GTPases Bind Raptor and Mediate Amino Acid Signaling to mTORC1. Science 320, 1496– 1501 (2008).

59. J. Lin, A. D. Hoppe, Uniform Total Internal Reflection Fluorescence Illumination Enables Live Cell Fluorescence Resonance Energy Transfer Microscopy. Microscopy and Microanalysis 19, 350–359 (2013).

60. J. Schindelin, et al., Fiji: an open-source platform for biological-image analysis. Nat Methods 9, 676–682 (2012).

61. M.-A. Bray, A. E. Carpenter, CellProfiler Tracer: exploring and validating high-throughput, time-lapse microscopy image data. BMC Bioinformatics, 1–7 (2015).

62. S. E. Quinn, et al., The structural dynamics of macropinosome formation and PI3-kinase-mediated sealing revealed by lattice light sheet microscopy. bioRxiv 94, 2020.12.01.390195 (2020).

63. T. Lambert, L. S., VolkerH. (2019)., tlambert03/LLSpy: v0.4.8 (Version 0.4.8) (2019).

