## Supplementary Figures for "IgG surface mobility promotes antibody dependent cellular phagocytosis by Syk and Arp2/3 mediated reorganization of Fcγ receptors in macrophages"

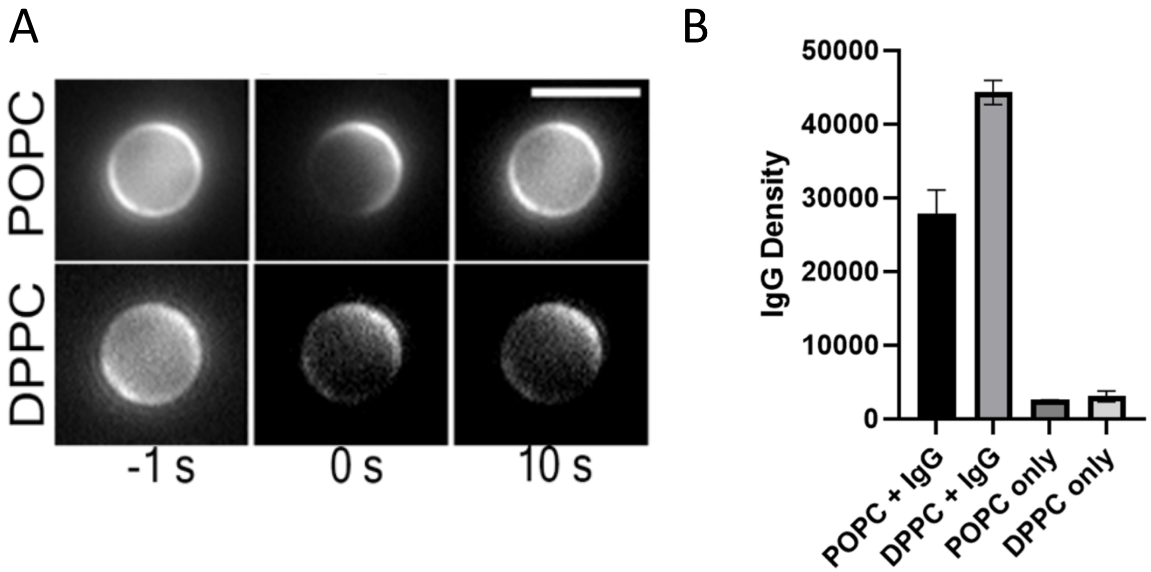


**Supplemental figure 1. IgG mobility and density on SLB-SBs.** (**A**) The mobility of anti-biotin-IgG2a-AF647 on SLB-SBs was assessed by Fluorescence Recover After Photobleaching. Mobile/fluid POPC SLB-SBs displayed free diffusion with uniform fluorescence recovery over the surface of the bead within 10 seconds. Gel-phase/immobile DPPC SLB-SBs did not allow redistribution of the IgG2a-AF647 (scale bar = 5 µm). (**B**) The relative density of IgG2a on SLB-SBs determined by flow cytometry using Alexa Fluor 633 Goat anti-Mouse IgG2a Cross-Adsorbed Secondary Antibody (ThermoFisher Scientific).

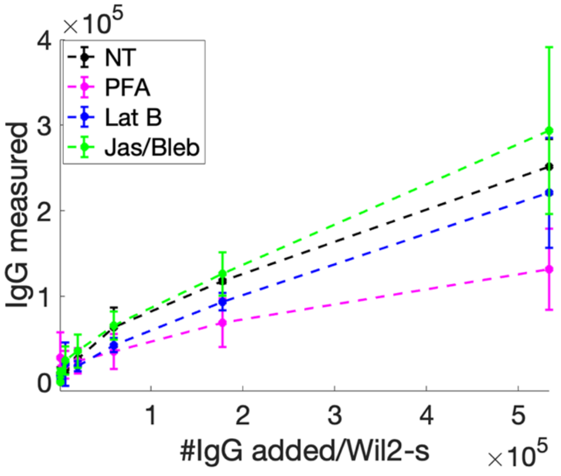


**Supplemental figure 2. Quantification of RTX density on WIL2-S cells.** RTX density on WIL2-S cell was measured as a function of the added number of RTX/WIL2-S cell using Alexa Fluor 633 Goat anti-Mouse IgG2a by flow cytometry. Geometric means and standard deviations are shown.

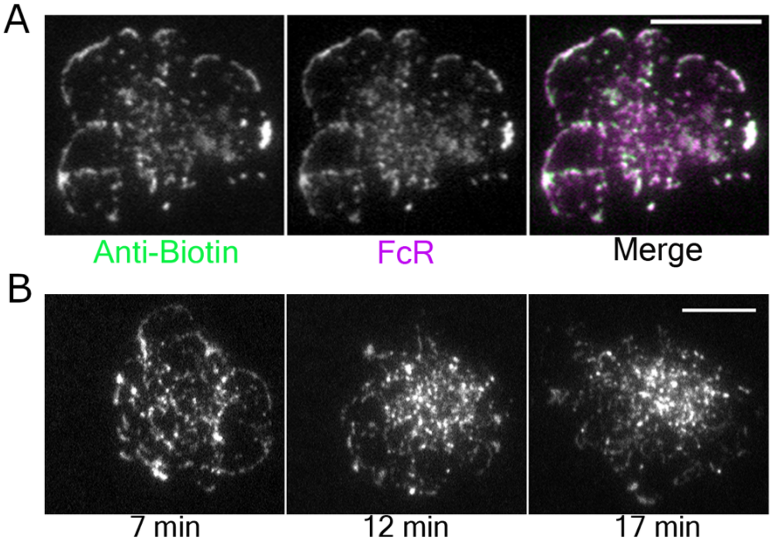


**Supplemental figure 3. IgG-FcγRI dynamics on SLBs.** (**A**) FcγRI-mScarlet was expressed in FLMs engaging POPC SLBs displaying IgG2a-AF647. Strong colocalization in microclusters indicates that the reorganization of IgG is the result of it's engagement by FcγR. (**B**) TIRF imaging on later time points of an FLM engaging on mobile/fluid POPC SLBs labeled with IgG2a-AF647 shows the aggregation of microclusters into patches that accumulate at the center of the contact site (scale bars = 10 μm).

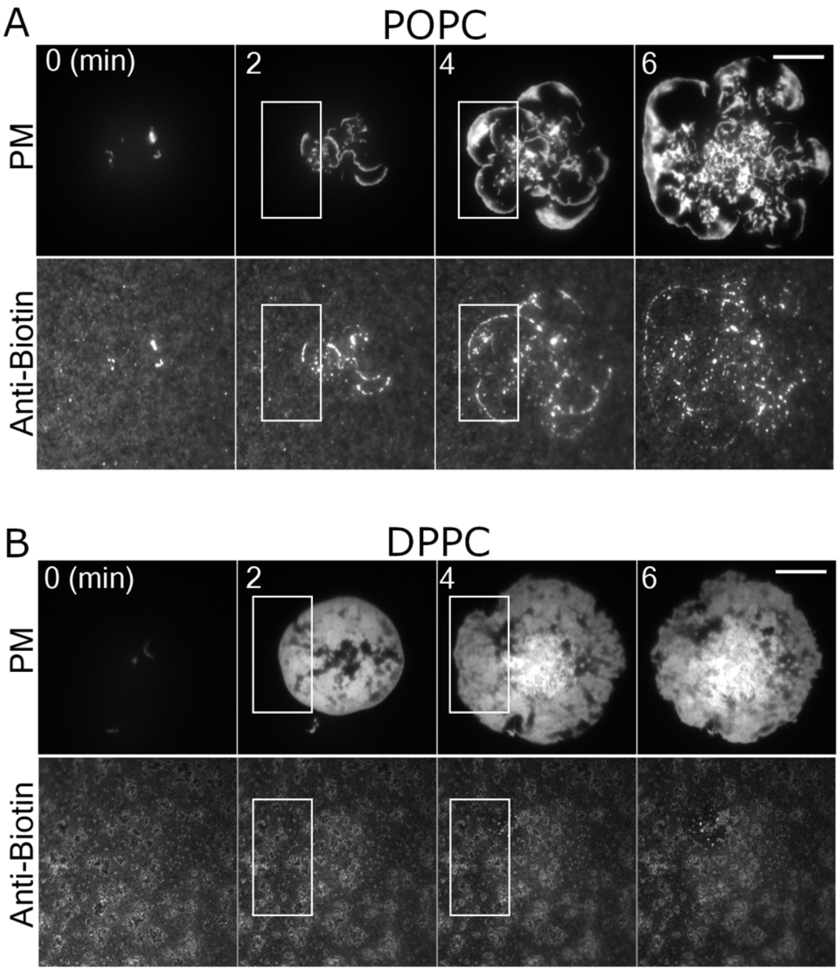


**Supplemental figure 4. IgG microcluster dynamics on POPC and DPPC SLBs.** TIRF imaging on an FLMs labeled with DiI engaging IgG2a-AF647 on mobile POPC SLBs (**A**) and immobile/gel-phase DPPC SLBs (**B**). Scale bars = 10 µm.
